# Single molecule mass photometry of nucleic acids

**DOI:** 10.1101/2020.01.14.904755

**Authors:** Yiwen Li, Weston B. Struwe, Philipp Kukura

## Abstract

Mass photometry is a recently developed methodology capable of detection, imaging and mass measurement of individual proteins under solution conditions. Here, we show that this approach is equally applicable to nucleic acids, enabling their facile, rapid and accurate detection and quantification using sub-picomoles of sample. The ability to count individual molecules directly measures relative concentrations in complex mixtures without need for separation. Using a dsDNA ladder, we find a linear relationship between the number of bases per molecule and the associated imaging contrast for up to 1200 bp, enabling us to quantify dsDNA length with 4 bp accuracy. These results introduce mass photometry as an accurate and rapid single molecule method complementary to existing DNA characterisation techniques.

## INTRODUCTION

Single molecule analysis has had a tremendous impact on our ability to study DNA structure, function and interactions (1). Next generation sequencing heavily relies on single-molecule methods, be it using single molecule fluorescence (2, 3) or nanopore-based approaches (4, 5). Similarly, single molecule methods are now heavily used in a variety of incarnations to study DNA-protein interactions (6), with both DNA and proteins visualised by fluorescence labelling to reach single molecule sensitivity (7). Label-free detection and quantification would be highly desirable in this context due to the associated reduction in experimental complexity and minimisation of potential perturbations caused by the sample itself. While visualisation of single DNA molecules has been possible for decades using non-optical methods, such as electron microscopy (8) and atomic force microscopy (9), which can also be used to study mechanical properties (10), label-free optical detection has remained a considerable challenge.

Label-free detection of single proteins has been reported for the first time in 2014 (11, 12) in the context of increasing sensitivity of interferometric scattering microscopy (13, 14). Further improvements to the detection methodology (15), recently lead to the development of mass photometry (MP), originally introduced as interferometric scattering mass spectrometry (16), which enables not only label-free detection and imaging of single molecules, but critically also their quantification through mass measurement with high levels of accuracy, precision and resolution at a lower detection limit on the order of 40 kDa. Given that biomolecules have broadly comparable optical properties in the visible range of the electromagnetic spectrum (17, 18), we therefore set out to investigate in this work to which degree the capabilities of MP translate to nucleic acids, which would enable not only their detection, imaging and analysis, but also provide a universal route to studying protein-DNA interactions at the single molecule level.

The operating principle behind MP is based on accurately measuring the change in reflectivity of a glass-water interface caused by interference between light scattered by a molecule binding to the interface and light reflected by that interface (**Fig. 1**). The experiment involves placing a small droplet of solution on top of microscope coverglass, to which molecules bind non-specifically, although in the case of DNA appropriate charging of the glass surface is advantageous to achieve tight binding (see Methods). We then visualise individual binding events on top of the static imaging background caused by residual substrate roughness by computing the differences between batches of averaged reflectivity images, which leads to the appearance and disappearance of single molecule signals from irreversible binding events in a continuous recording (**Supplementary Movie 1**) (12, 15). By determining the point in time when each individual molecule binds, we can then quantify the associated reflectivity change, yielding highly accurate, precise and resolved contrast distributions.

**Figure 1.**
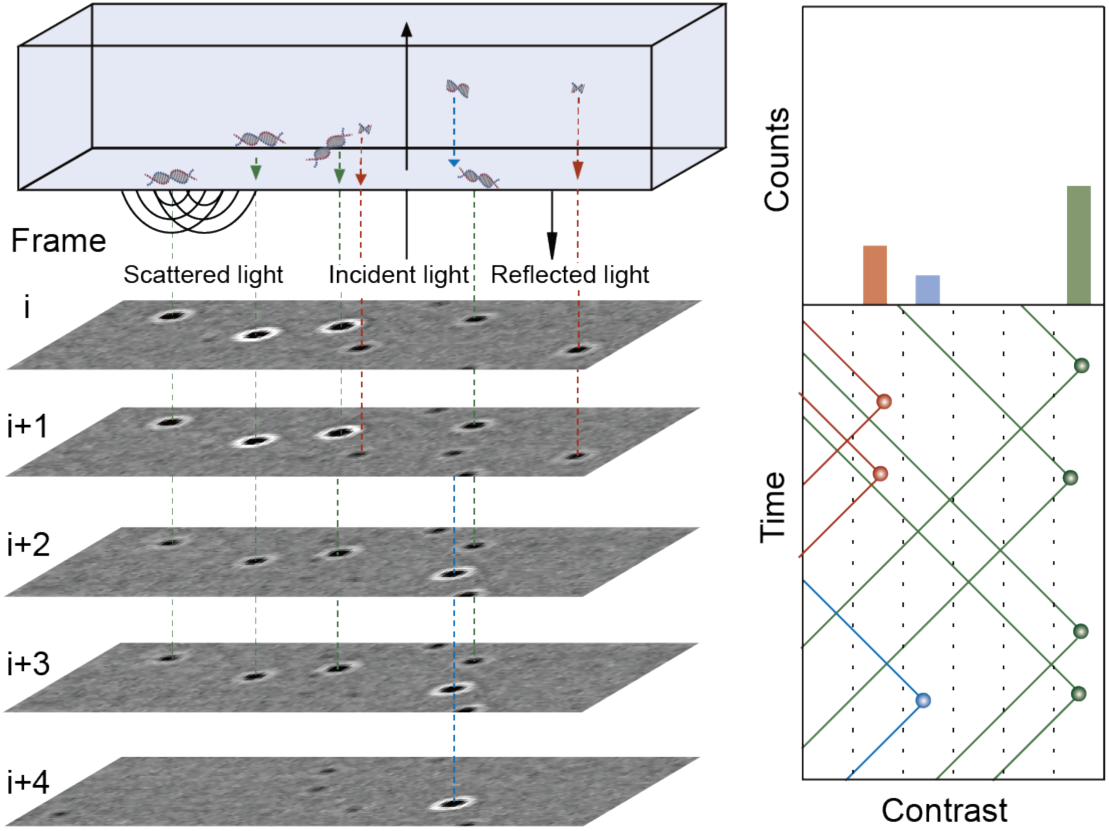
Working principle of label-free DNA detection and quantification by mass photometry. Individual DNA molecules diffusing in solution bind to an appropriately charged glass surface. Individual binding events cause changes to the reflectivity of the interface, visualised by a contrast-enhanced interferometric scattering microscope through the interference between scattered and reflected light. The need to remove the static imaging background requires a differential imaging scheme, resulting in irreversible binding events appearing as signals that increase and decrease in time. The single molecule signal corresponds to the maximum observed signal, reaching a maximum at different time points depending on the arrival time of the specific species.

## MATERIALS AND METHODS

Solvents and chemicals were from Sigma Aldrich unless otherwise noted. Milli-Q water and high-grade solvents were used for all experiments. A double-stranded DNA ladder consisting of 100, 200, 400, 800, 1200 and 2000 base pairs was purchased from Invitrogen (Cat. No. 10068013). Single stranded DNA with 4536, 6048, 7249 and 8064 bases was prepared as previously described (19). Single stranded DNA (155-mer) was synthesised on an Applied Biosystems 394 automated DNA/RNA synthesiser using a standard 0.2 μmole phosphoramidite cycle of acid-catalysed detritylation, coupling, capping, and iodine oxidation. Stepwise coupling efficiencies and overall yields were determined by the automated trityl cation conductivity monitoring facility and was >98.0%. Standard DNA phosphoramidites and additional reagents were purchased from Link Technologies Ltd, Sigma-Aldrich, Glen research and Applied Biosystems Ltd. All beta-cyanoethyl phosphoramidite monomers were dissolved in anhydrous acetonitrile to a concentration of 0.1 M immediately prior to use with a coupling time of 50 s. Cleavage and deprotection were achieved by exposure to concentrated aqueous ammonia solution for 60 min at room temperature followed by heating in a sealed tube for 5 h at 55 °C. Purification was carried out by denaturing 8% polyacrylamide gel electrophoresis. In brief, Formamide (500 µL) was added to the DNA sample (500 µL in water) before loading to the gel, bands corresponding to the full length were excised and the DNA was isolated using the ‘crush and soak method’. The excised polyacrylamide pieces were broken down into small pieces then suspended in distilled water (25 mL). The suspension was shaken at 37 °C for 18 h then filtered through a plug of cotton wool. The filtrate was concentrated to approximately 2 mL then desalted using two NAP-25 followed by one NAP-10 columns. The desalted eluent was lyophilised prior to use.

Prior to MP measurements, 231 nM (0.1175 µg/µl) double-stranded DNA stock solutions were diluted 25-fold in 5 mM Tris, 10 mM MgCl_2_, pH = 8. Single-stranded DNA stock solutions (167 nM for 4536 bp, 125 nM for 6038 bp, 100 nM for both 7249 bp and 8064 bp) were diluted 10-fold in the same buffer. 7.79 µM 155 bp single-stranded DNA stock solutions were diluted 1000-fold. Standard protein marker solutions were diluted 10-fold in the same buffer. Samples were kept at room temperature during analysis.

### Mass photometry

Microscope coverglass (24 × 50mm # 1, 5 SPEZIAL, Menzel-Glaser) and APTES-functionalised coverslips were prepared as described previously (16, 20). Briefly, coverslips were cleaned by sequential sonication in 2% Hellmanex (Hellma Analytics), water and iso-propanol for 10 minutes before plasma cleaning with oxygen (Diener electronic Zepto) for 8 minutes. The coverslips were then immersed in 200 ml 2% APTES solution in acetone for 1 minute with agitation before rinsing in 200 ml acetone. Finally, the coverslips were incubated at 110°C for one hour and cleaned by sonication in isopropanol (10 minutes) and water (5 minutes) before drying under a nitrogen stream.

Mass photometry was performed using a home-built microscope described elsewhere (15, 16). Instrument settings were as follows: Laser wavelength: 520 nm, Laser power: 300 mW, frame rate = 955 Hz, exposure time = 998 µs, temporal averaging: 5-fold, pixel binning: 4×4, field of view: 3.5 × 10 µm. This leads to an effective frame rate of 191 Hz and a pixel size of 84.4 nm prior to further analysis. All measurements were performed using flow chambers made by microscope cover glass and double-sided tape with 15 µl sample per analysis. Flow chambers were first filled with a buffer blank to position the coverslip into the optimal focus position. Samples were then added to one side of the flow chamber and introduced by capillary flow with the aid of tissue paper to draw liquid into the chamber. Data acquisition was started within 15 seconds of sample addition for a total of 120 seconds. In total, 5 replicates were taken for the double-stranded DNA ladder and 3 replicates for each single-stranded DNA sample. Data acquisition was performed using custom software written in LabView, generating a single movie file (.tdms) for further analysis.

### Data analysis

All acquired movies were processed and analysed using Discover MP v1.2.4 (Refeyn Ltd). The analysis procedure involved two fitting parameters for identifying landing events: 1) Threshold 1 related to a given particle contrast amplitude relative to the background; and 2) Threshold 2 related to the radial symmetry of the detected point spread function (PSF) of the same particle.

Analysis parameters for dsDNA and ssDNA samples are shown in Table 1.

| DNA sample | Number of binned frames, $n$ | Threshold 1 | Threshold 2 |
| --- | --- | --- | --- |
| dsDNA ladder | 8 | 1 | 0.2 |
| ssDNA 155bp | 8 | 1 | 0.15 |
| ssDNA 4536bp | 5 | 3 | 0.3 |
| ssDNA 6048bp | 5 | 3 | 0.3 |
| ssDNA 7249bp | 5 | 3 | 0.3 |
| ssDNA 8064bp | 5 | 3 | 0.3 |

The output h5-files contained a list of all detected particles within the analysed movie and their corresponding contrast values. The contrasts of all landing events were plotted as a scatter plot along the time axis. A histogram of the number of landing events and the contrasts was then generated. The resulting peaks were fit to a sum of Gaussians and the mean of the fitted peaks was taken as the contrast for each DNA component. The base pair to contrast ratio was determined by a linear fit. Base pair error was given as a deviation of the measured number of base pairs from the nominal number given by the manufacturer.

### Diffusion Correction and Concentration Measurement

The relative abundance of each DNA fragment in the dsDNA ladder was calculated from the area of each Gaussian peak in the kernel density estimate (KDE) plot, 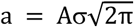, where *a* is the area, *A* is the amplitude and *σ* is the standard deviation of the fitted Gaussian. The contrast magnitude achievable with the 100 bp species approached the detection limit of the instrument. As a result, while events could be clearly detected, the detection fidelity was imperfect leading to variations in the number of detected events as a function of analysis parameters.

To account for differences in binding rate and thus molecule counts caused by differences in diffusion coefficient, we applied a correction to the measured mass distributions (16). We assumed that the binding rate constant scales with the diffusion coefficient, which has been reported to be roughly proportional to (base pair)^−0.72^ for DNA (i.e., *k*_*i*_ = α * *bp*^−0.72^, where *k*_*i*_ is the binding rate constant for DNA component *i* and *α* is a scaling factor) (21). We assumed that the scaling factor *α* is constant for all DNA components. To estimate the scaling factor *α*, an exponential function was fitted to the number of landing events *vs* time and to obtain an average binding rate, *k*. The scaling factor was calculated as: 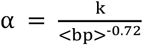, where <bp> is the average number of base pairs of all the DNA components in solution calculated based on the distribution of each DNA component, 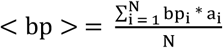, where *bp*_*i*_ is the number of base pairs and *a*_*i*_ the relative abundance measured experimentally of DNA component *I*, and *N* is the total number of species in the solution. To accurately estimate the proportion of each DNA fragment present in solution, it is important to account for landing events that occur between the time when the sample is added (*t*_*addition*_) and when data acquisition starts (*t*_*0*_). This was solved by fitting the exponential decay of measured binding events (from t_addition_) from the addition of sample to completion of all sample binding (*t* = infinity). Experimentally, we integrated from a given time *t*_*0*_ = 15 s after addition of sample up to a later time, *t*_*final*_ = 135 s, when the acquired movie ended. Relating these two, the corrected intensity was given by: 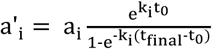, where *a’*_*i*_ is the intensity of DNA component *i* corrected over all time, and *a*_*i*_ is the experimentally measured abundance. The corrected mass distribution was then renormalized.

## RESULTS

To test the applicability of MP to a representative DNA sample, we begin with an analysis of a standard low mass dsDNA ladder. A 9 nM solution led to distinct molecular binding events with clearly varying molecule-to-molecule contrasts (**Supplementary Movie 1**), while significant unbinding events were observed on non-APTES coverslips (**Supplementary Movie 2**), suggesting that appropriate surface charge is required to achieve clear binding events. The contrast histogram of the landing events on non-APTES coverslips showed very poor resolution (**Supplementary Movie 2B**). As for signals collected on APTES coverslips, a scatter plot of these signals obtained by quantifying the signal magnitude for each individual binding event exhibited six clear bands, as expected from the ladder used (**Fig. 2**). This separation persisted upon binning into a contrast histogram, with baseline resolution from the second peak onwards. The observed spacing and knowledge of the ladder composition allowed for assignment to different contour lengths by inspection. These results demonstrate that MP can detect single DNA molecules without labels with a comparable performance in terms of mass sensitivity and resolution to polypeptides.

**Figure 2.**
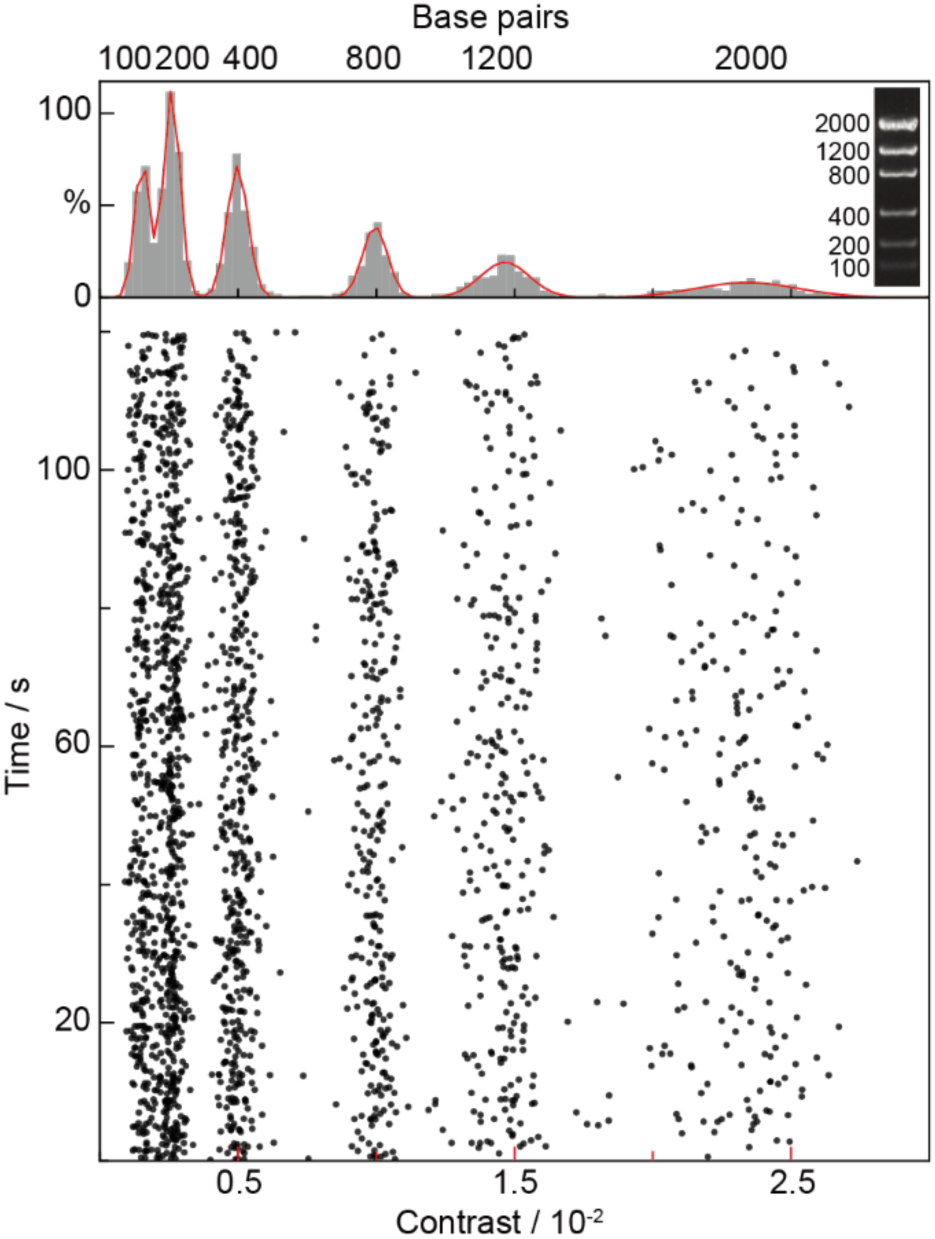
Scatter plot and resulting contrast histogram obtained by quantifying the image contrast on a molecule-by-molecule basis for a low mass dsDNA ladder (see inset). The close correspondence between the gel and resolvable features in the contrast histogram allows for assignment by inspection.

Quantifying the collision frequency should provide direct information on molecular concentration for each species assuming label-free, universal detection of all binding events. Multiple repeats of the ladder experiment exhibited high reproducibility (7.7% RMS) in the total number of detected molecules, despite the simplicity and inherent variability of the measurement due to manual sample addition and timing when recording is started (**Fig. 3a**). The relative fluctuations between the peak areas amounted to 12 ± 3.3% RMS (**Fig. 3b, blue dots**). Despite the fact that the ladder contains an equimolar mixture of molecules, we observed clear variations in peak areas with a drop towards larger species. The collision frequency of molecules with the surface, and thus detection rate, however, is not only a function of solution concentration, but also diffusion coefficient, which decreases considerably with contour length (21). Our qualitative observation of a decrease in binding events per measurement with contour length agrees with this expectation.

**Figure 3.**
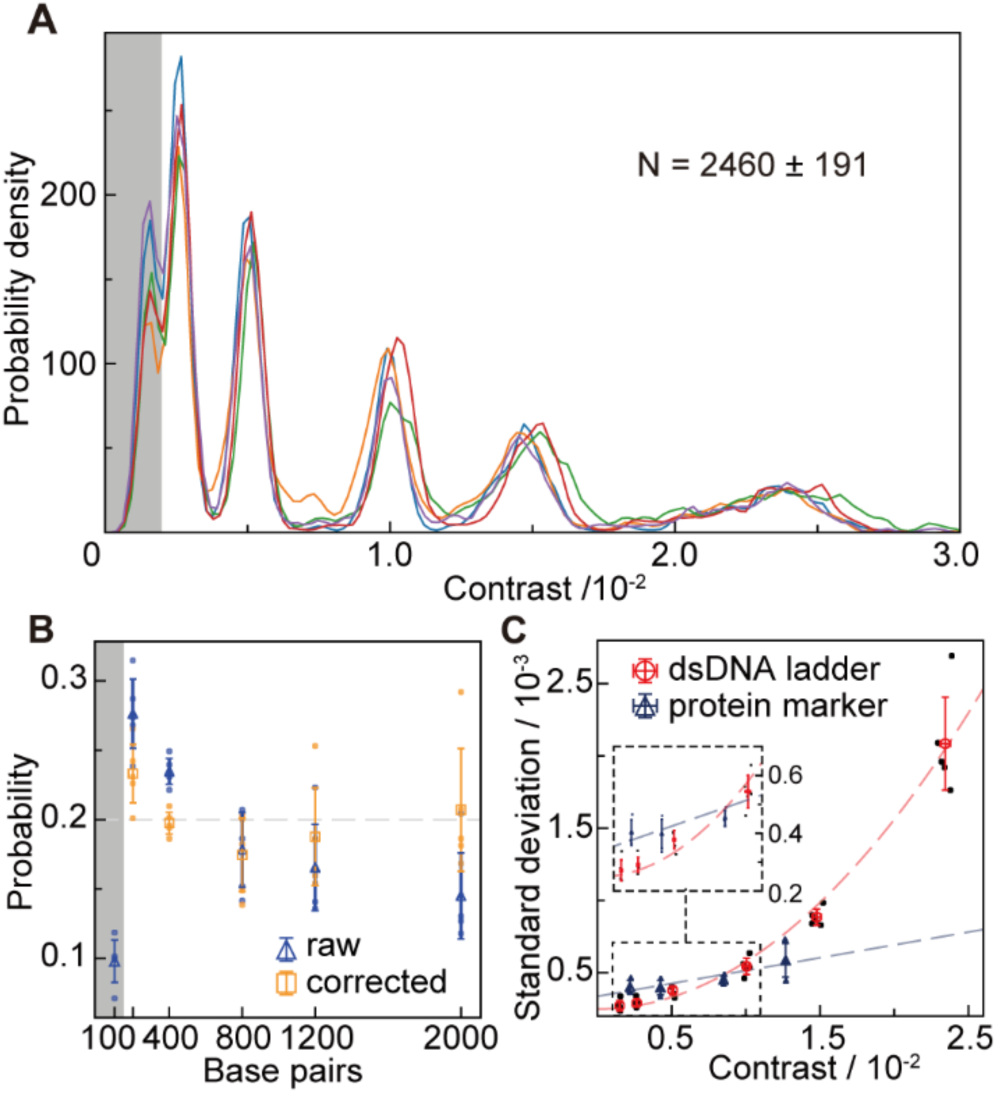
Achievable concentration precision and base pair resolution. **A**, Reproducibility of individual MP measurements of the same dsDNA ladder sample. The plots were generated from mass histograms using a Kernel Density of width 2.1 × 10^−4^. **B**, Extracted mole fractions before and after correction for length-dependent diffusion. **C**, Comparison of contrast resolution between dsDNA and a globular protein mixture of comparable imaging contrast for the same instrument.

To account for these differences, we need to relate the number of molecules that have bound during our finite measurement window to the number we would have observed for an infinite observation time where all molecules would be depleted from solution by surface binding. Since smaller molecules diffuse more quickly, more of them will be removed from solution initially, resulting in a concentration difference once the measurement is started compared to the original solution (16). Correcting for this behaviour normally increases the amount of large relative to small molecules (**Fig. 3b, orange dots**), in this case resulting in effectively equimolar concentrations for all species of contour length 200 bp or larger. The lower than expected concentration of 100 bp species was most likely caused by non-unity detection efficiency as this species approaches the detection limit of MP in the implementation used here. Nevertheless, these results demonstrate the ability of MP to quantify relative amounts of DNA in solution above 100 bp simultaneously without need for separation.

Careful inspection of the obtained mass distribution in Fig. 2a reveals a clear variation in peak width with molecular size, with a lower width for small species and much larger widths for large species compared to globular proteins producing similar imaging contrast (**Fig. 3c**). These results point towards a variability in contrast for dsDNA possibly due to binding events occurring with different conformations, and thus effective density and polarizability, which determines the magnitude of the detected signal in an interferometric measurement (14). We would expect this effect to become relatively more pronounced the longer the DNA molecules are, given the increased structural flexibility in the light of the persistence length of DNA (∼150 bp), in line with our observations. Similarly, the reduced width for small species (<400 bp) can be explained by a comparatively lower degree of disorder in terms of structure and thus polarizability given the structural rigidity of DNA on short length scales compared to globular proteins binding non-specifically to a glass surface.

The observed peak spacing roughly matches the spacing expected for a direct proportionality between the MP contrast and the number of base pairs. To quantify this correlation, we repeated these measurements 5 times, finding almost perfect correspondence (R^2^ = 0.9998 ± 0.0001) for all species up to 1200 bp, with a slightly lower than expected contrast for the largest (2000 bp) species (**Fig. 4a, left**). The resulting conversion from imaging contrast to contour length amounts to 1.22 ± 0.02×10^−5^ /bp for dsDNA. Applying this conversion factor to each individual measurement allowed us to determine the average base pair error, which amounted to 3.9 ± 4.3 bp up to 1200 base pairs, with slight variations as a function of molecular size (**Fig. 4b**). At this stage it is unclear to which degree the observed error is indeed representative of the limits achievable by MP or whether they are caused by sequence-specific variations in molecular mass or molecular polarizability, which we could not account for given that the sequence of the ladder components were unknown.

**Figure 4.**
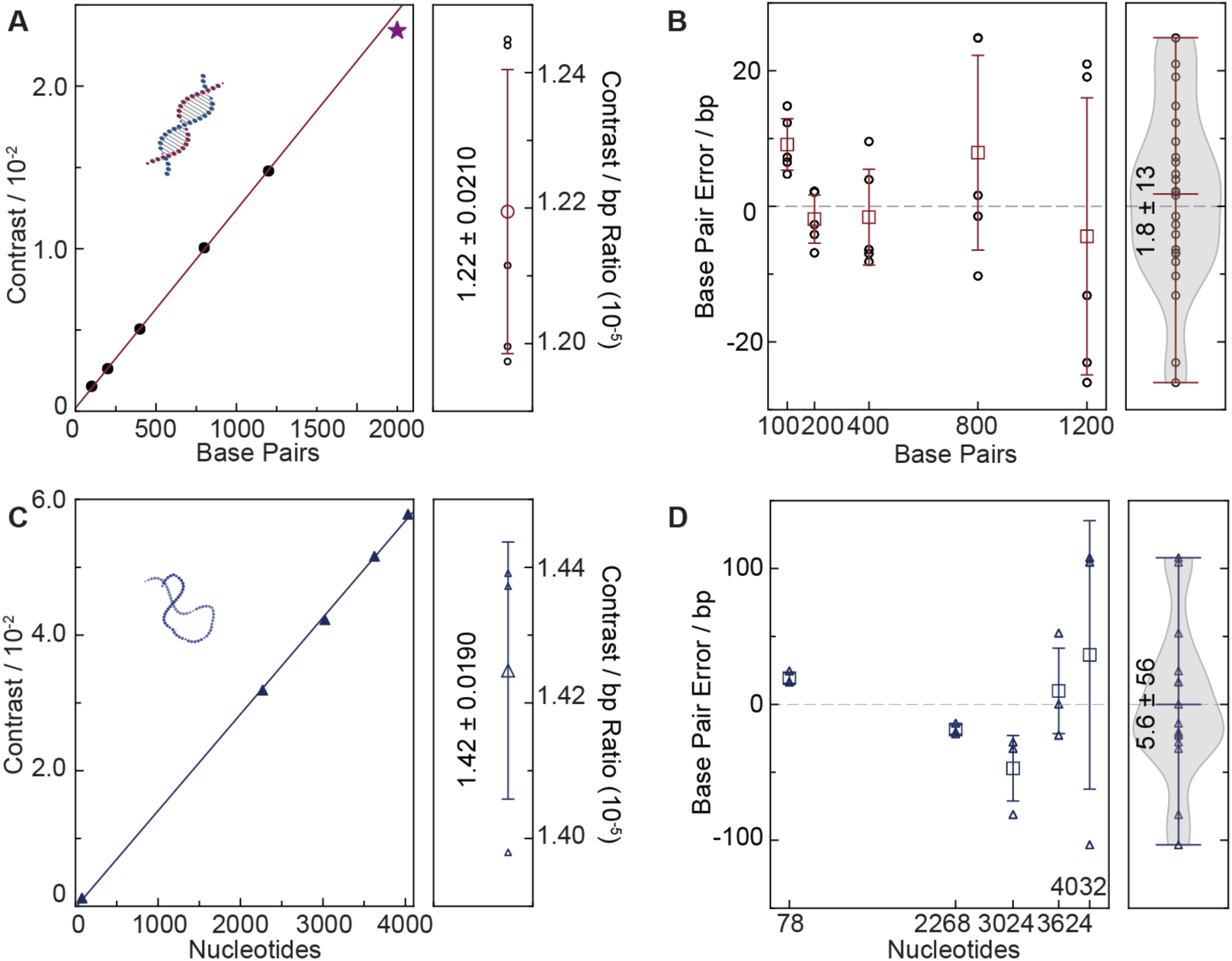
Characterisation of nucleotide accuracy and precision for dsDNA and ssDNA. **A**, Correlation between imaging contrast and base pair number. The 2000 bp data point (star) was omitted for the calibration due to the molecular size becoming comparable to the diffraction limit (200 nm). **B**, Resulting base pair accuracy for independent measurements using the average contrast-to-bp conversion. **C**,**D**, Equivalent measurements for ssDNA. The bandwidths for the violin plots are 0.32.

Repeating the same process with ssDNA revealed a similarly linear relationship between the number of bases and the imaging contrast, without noticeable deviations for larger species (**Fig. 4c**). The associated average basepair error of 23 ± 15 bp is poorer (**Fig. 4d**), although this is may have been partially caused by the need to run separate experiments for each species, rather than mixtures as for dsDNA due to insufficient sample purity. The contrast-to-basepair ratio is slightly larger for ssDNA at 1.42 ± 0.0190×10^−5^ /bp, likely due to a higher effective density of ssDNA in the presence of Mg^2+^ due to its increased flexibility or increased polarizability in the absence of basepair hybridisation (22). The lack of non-linear contrast behaviour even for very long ssDNA molecules is likely due to the fact that ssDNA is highly compacted under the buffer conditions used, ensuring essentially uniform base pair densities for all studied species irrespective of the number of base pairs in contrast to dsDNA, where the contour length of DNA plays a non-negligible role.

## DISCUSSION

Taken together, these results establish mass photometry as a novel analytical approach to studying nucleic acids, with a range of potential future analytical applications. We have demonstrated both absolute and relative concentration measurements based on molecular counting with comparable precision to UV-absorption based approaches, but with the specific advantage of operation at low concentrations (nM) and minimal sample requirements, currently only limited by our sample delivery approach. Given the field of view, this could be reduced to (100µm)^3^ in the future, allowing detection and quantification of as little as attograms of DNA. The current base pair accuracy of 4 bp is comparable to unreferenced CE, and can be improved further by using appropriate internal standards or possibly even through precise knowledge of the calibrant sequence in the future. Base pair resolution will likely never reach that achievable with electrophoretic methods, but the solution operation of MP provides much potential for combination with such approaches in the future. Given the single molecule nature of MP, coupled with its intrinsic compatibility for visualising and quantifying proteins, and suitability for combination with single molecule fluorescence imaging (23), MP is likely to become an powerful addition to the existing toolbox single molecule methodologies aimed at quantifying and studying nucleic acids and their interactions.

## ACKNOWLEDGEMENT

We thank the Brown group (University of Oxford) for providing the 155mer of ssDNA and the Dietz Group (TU Munich) for the ssDNA scaffolds.

## CONFLIC OF INTEREST

PK is academic founder and director of Refeyn Ltd.

