## Supplementary for "Single molecule mass photometry of nucleic acids"

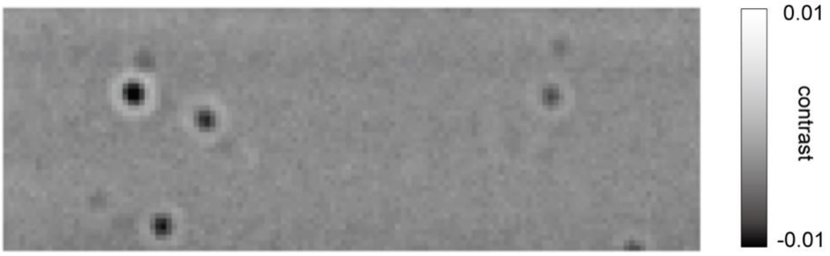


**Supplementary Movie 1:** Ratiometric movie of the dsDNA ladder binding to an APTES functionalised coverslip.

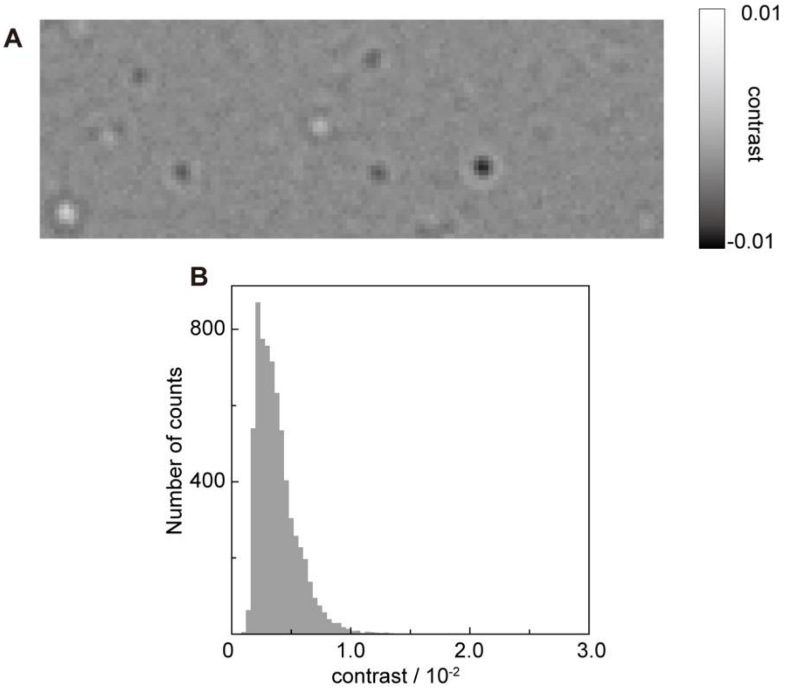


**Supplementary Movie 2: A.** Ratiometric movie of dsDNA ladder binding to a regular glass coverslip. **B.** Corresponding histogram of the landing events
